# Φ-Space: Continuous phenotyping of single-cell multi-omics data

**DOI:** 10.1101/2024.06.19.599787

**Authors:** Jiadong Mao, Yidi Deng, Kim-Anh Lê Cao

## Abstract

Single-cell multi-omics technologies have empowered increasingly refined characterisation of the heterogeneity of cell populations. Automated cell type annotation methods have been developed to transfer cell type labels from well-annotated reference datasets to emerging query datasets. However, these methods suffer from some common caveats, including the failure to characterise transitional and novel cell states, sensitivity to batch effects and under-utilisation of phenotypic information other than cell types (e.g. sample source and disease conditions).

We developed Φ-Space, a computational framework for the continuous phenotyping of single-cell multi-omics data. In Φ-Space we adopt a highly versatile modelling strategy to continuously characterise query cell identity in a low-dimensional phenotype space, defined by reference phenotypes. The phenotype space embedding enables various downstream analyses, including insightful visualisations, clustering and cell type labelling.

We demonstrate through three case studies that Φ-Space (i) characterises developing and out-of-reference cell states; (ii) is robust against batch effects in both reference and query; (iii) adapts to annotation tasks involving multiple omics types; (iv) overcomes technical differences between reference and query.

The versatility of Φ-Space makes it applicable to a wide range analytical tasks beyond cell type transfer, and its ability to model complex phenotypic variation will facilitate biological discoveries from different omics types.

## 1 Introduction

Single-cell multi-omics technologies have revolutionised molecular cell biology by providing multi-omic measurement of cells, including genome, transcriptome, epigenome and proteome. These technologies enable an increasingly refined definition of cell states, which deepens our understanding of cellular heterogeneity and biological complexity [1]. However, the fast generation of population-level and atlas-scale single-cell data has created a huge demand for computationally efficient and statistically reliable annotation of cell populations in emerging datasets [2, 3]. Classifying cells into cell types or states, an analytical task referred to as cell annotation, has been identified as one of the ‘grand challenges’ in single-cell data science [4].

There are two major approaches for cell type annotation: *de novo* annotation based on marker genes and automated annotation based on reference datasets [4–6].

The *de novo* approach is well established, and consists in applying unsupervised clustering to identify homogeneous cell groups, before manually annotating each group based on their (known) marker genes. However, this approach suffers from two major drawbacks: (i) it is labour-intensive and hence lacks scalability; and (ii) it is based on the selection of marker genes and cell type labels that are somewhat arbitrary. These drawbacks result in a lack of scalability and reproducibility [4]. In addition, assigning to a cell cluster a unique cell type label is not appropriate for cells undergoing continuous transition, since discrete clustering often fails to capture within-cluster heterogeneity of developing cells [7].

Automated annotation computationally transfers cell type information from high-quality annotated *reference* datasets to *query* datasets [5]. This approach often uses supervised classification: a classification model (e.g. *k*-nearest neighbour, random forest or deep neural network) is trained on the reference dataset then assigns a cell type label to each cell in the query single-cell dataset [6, 8–12]. Annotation based on supervised classification has first been developed within omics [13], e.g. scRNA-seq reference and query. Methods for querying scRNA-seq data against bulk references have also been developed to leverage the rich phenotypic information contained in bulk reference atlases [14, 15]. More recently, Hao et al. [16] developed a cross-omics annotation method, where the reference and the query consist of features of different omics types, e.g. scRNA-seq reference and scATAC-seq query.

However, current annotation methods suffer from some common caveats. First, existing methods mainly view cell type annotation as a *hard classification* problem, focusing on accurately predicting the cell types of query cells. These methods lack the power to characterise continuous and transitional cell states. Although methods such as SingleR [12], Seurat V3 [9] and Celltypist [11] do provide some continuous quantification of the predicted cell types, these *soft classification* results remain under-utilised. Second, existing methods assign to each cell only one label, so they are unable to jointly model multiple layers of phenotypes (e.g. cell type and sample source). Third, few existing methods (except Aran et al. [12]) can directly use bulk data as reference, due to the much higher proportion of zero counts in scRNA-seq data compared to bulk [15]. As a result, most methods cannot utilise the rich phenotypic information in population-level bulk atlases such as the bulk atlas generated by Angel et al. [14]. Fourth, a cell atlas usually consists of data from multiple experimental batches and studies, and hence suffer from strong and complex batch effects. This poses additional computational challenges for existing annotation methods, since they require the appropriate correction of batch effects within the reference [17]. Lastly, existing methods lack the flexibility to be extended to multi-omics data, resulting in a under-utilisation of the increasingly common single-cell multi-omics references [17].

To overcome these limitations, we have developed Φ-Space that uses soft classification to recover the continuous nature of cell states and then use the annotation results as input to downstream analyses. The main innovation of Φ-Space is the phenotype space analysis modelling strategy: the key idea, inspired by Delaigle and Hall [18], is to view soft classification as a dimension reduction to phenotype space and then conduct downstream analyses therein. More specifically, in Φ-Space we assign to each query cell a membership score on a continuous scale for each reference phenotype. Thus each query cell is continuously characterised in a multi-dimensional *phenotype space*. The phenotype space enables various downstream analyses, including insightful visualisations, clustering but also hard classification.

Compared to analyses in the original omics space, which is the space defined by the original omics features such as gene or protein expressions, we demonstrate that phenotype space analysis is robust against batch effects in both the reference and the query data. The phenotype space analysis makes Φ-Space much more flexible than conventional annotation methods. In particular, Φ-Space can jointly model multiple layers of phenotypes in both bulk and single-cell references. In addition to within- and cross-omics annotation, Φ-Space can also handles multi-omics annotation, where both reference and query contain multimodal measurements (e.g. CITE-seq, gene expression and surface protein). We show that Φ-Space is highly versatile, and its versatility does not come with a high computational price as Φ-Space is based on linear factor modelling using Partial Least Squares regression (PLS) [19]. Due to PLS’s ability to remove unwanted variation, no additional batch correction nor harmonisation of reference and query is needed.

The remainder of the manuscript is organised as follows. In Section 2 we describe the Φ-Space framework and three case study datasets ranging from transcriptomics (bulk and single-cell RNA-seq), proteomics (CITE-seq) to epigenomics (scATAC-seq). In Section 3 we showcase the versatile use of Φ-Space in uncovering phenotypic variations by continuous phenotyping in the case studies. We then conclude and discuss areas of future work in Section 4.

## 2 Materials and methods

### 2.1 Φ-Space continuous phenotyping

#### Notations

In the following sections, we denote the reference dataset by (*X*_ref_ *, Y*_ref_ ), where *X*_ref_ is a (*N ×P* ) omics feature matrix of normalised values, with *P* omics variables measured on *N* samples (either bulk or single-cell; depicted in Fig. 1A). The phenotype matrix *Y*_ref_ is a (*N × K*) dummy matrix that represents the *K* phenotype labels of the *N* samples, that is, each column of *Y*_ref_ represents a phenotype label (e.g. a cell type or sample source) and a sample is assigned the value 1 if it has that label or *−*1 otherwise. Importantly, we allow each sample to have more than one label (i.e. a sample can be both dendritic cell and *in vitro*), thus each row of the matrix *Y* may contain multiple values of 1 .

**Fig. 1.**
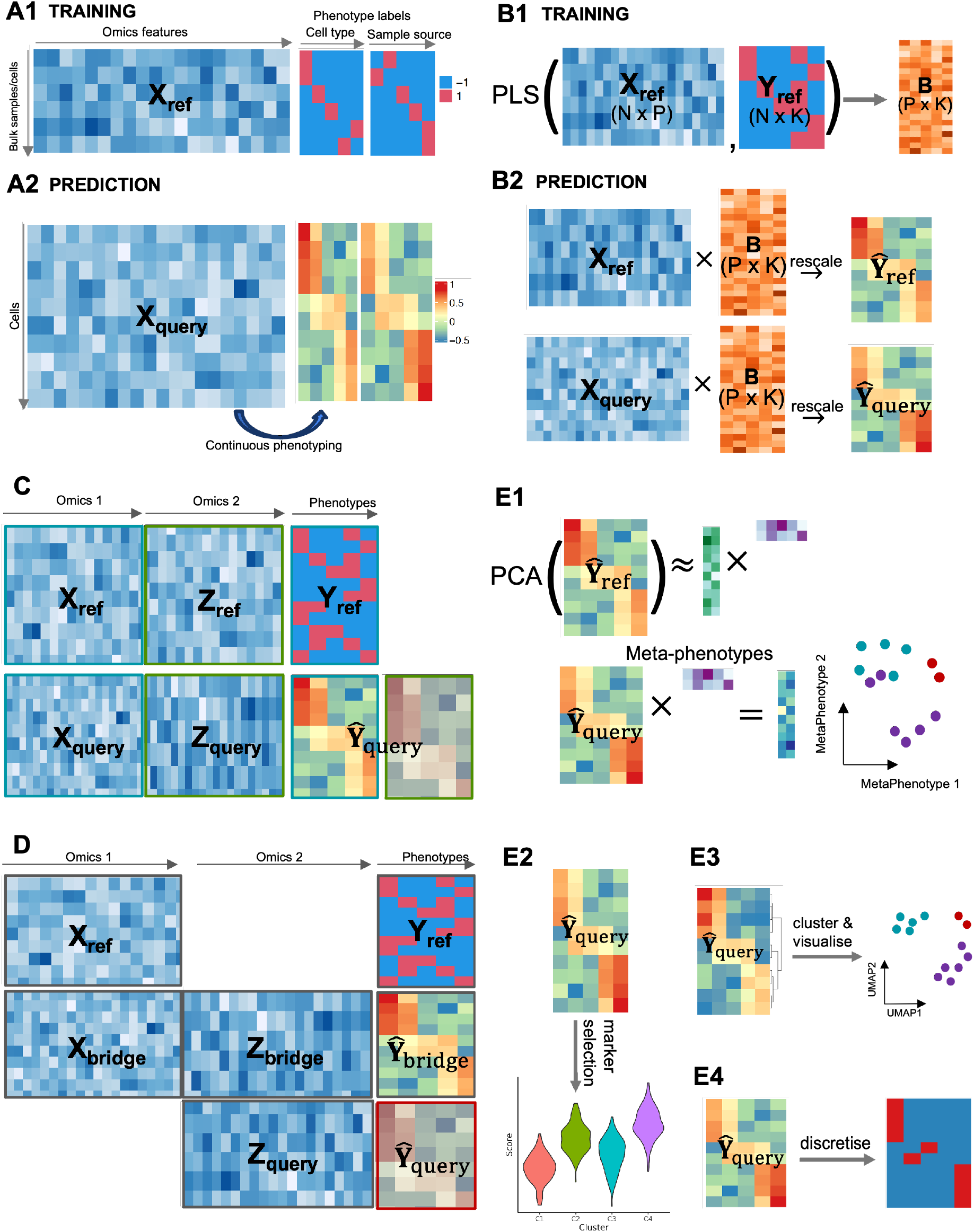
Overview of Φ-Space. **A**) Φ-Space continuous phenotyping based on **A1**) a reference dataset with discrete phenotype labels, e.g. cell types and sample sources; **A2**) the query cell states are then continuously characterised. B) The core method used in Φ-Space is partial least squares (PLS) regression. **B**1) For within-omics annotation, a PLS regression model is trained on the reference data (*X*_ref_ *, Y*_ref_ ) to compute *B*, a matrix converting omics features to continuous phenotype scores. **B2**) The phenotype space embeddings of both reference and query are computed as rescaled versions of *X*_ref_ *B* and *X*_query_*B*. **C**) We conduct multi-omics annotation when both the reference and the query data contain multiple matching modalities (different types of omics measurements for the same cells). We first conduct within-omics annotation for each modality independently, and then concatenate the phenotype space embeddings derived from different modalities. This multi-omic phenotype space embedding is then used for downstream analysis. **D**) We conduct cross-omics annotation when the reference *X*_ref_ and the query *Z*_query_ contain different omics features. We utilise a bimodal bridge dataset (*X*_bridge_*, Z*_bridge_) sharing omics features with both reference and query. We first annotate the bridge cells using the reference modality, i.e. computing *Ŷ*_bridge_ as rescaled *X*_bridge_*B*. We then train a new PLS model for the relationship between the query modality of bridge *Z*_bridge_ and the derived continuous annotation *Ŷ*_bridge_, resulting in *B*_bridge_. Finally we annotate the query data *Z*_query_ by computing *Ŷ*_query_ as rescaled *Z*_query_*B*_bridge_. **E**) We can conduct the following downstream analyses based on the phenotype space embeddings *Ŷ*_ref_ and *Ŷ*_query_. **E1**) Reference mapping: we reduce the dimension of *Ŷ*_ref_ by principal component analysis (PCA), where each PC is interpreted as a meta-phenotype in the reference, and then *Ŷ*_query_ are mapped to this PC space via the loading vectors learned from *Ŷ*_ref_ . **E2**) Marker selection: given some grouping of query cells (e.g. disease conditions, groups defined by clustering analysis), we can identify phenotypic markers (e.g. enriched cell types) of each group, which provides more interpretable and . **E3**) Clustering: we apply clustering algorithms to *Ŷ*_query_ to identify biological meaningful cell states in the query. **E4**) Hard classification: for each query cell, we select the highest scored cell type as the predicted cell type for that cell.

#### Training step

To model the relationship between the omics feature matrix *X*_ref_ and the multivariate *Y*_ref_ , we apply partial least squares regression (PLS) [19, 20] to estimate a low-rank regression coefficient matrix *B* of size (*P × K*) so that *Y*_ref_ *≈ X*_ref_ *B* (depicted in Fig. 1B). The rank of *B* is equal to the number of PLS components, by default set to *K*, the number of all phenotype labels. Supplementary Section S1 describes in detail how to optimise the selection of this parameter.

Feature selection is often desired to remove unwanted variation contained in noisy features and to increase the model’s interpretability. PLS regression allows for feature selection in *X*_ref_ so that only the top features that are highly predictive are selected in PLS; see Section S1 for details.

#### Annotation step

To transfer the phenotypic information learnt from the reference, we consider the three analytical tasks:

1. **Within-omics annotation** (Fig. 1B). When the query data *X*_query_ of size (*M × P* ), with *M* denoting the number of query cells, contain the same omics variables as *X*_ref_ , we compute the annotation denoted *Ŷ*_query_ based on *X*_query_ and the regression coefficient matrix *B* from PLS(*X*_ref_ *, Y*_ref_ ), as detailed in Supplementary Methods Section S1. We illustrate this analysis in the DC case study in Section 3.1.
2. **Multi-omics annotation** (Fig. 1C). When both the reference (*X*_ref_ *, Z*_ref_ ) and the query (*X*_query_*, Z*_query_) contain two modalities, we train PLS(*X*_ref_ *, Y*_ref_ ) and PLS(*Z*_ref_ *, Y*_ref_ ), and then concatenate the predicted scores to obtain the annotation *Ŷ*_query_. We illustrate this analysis in the CITE-seq case study in Section 3.2.
3. **Cross-omics annotation** (Fig. 1D). When the query data *Z*_query_ of size (*M × Q*), with *M* denoting the number of query cells and *Q* the number of omics variables, is of a different omics type compared to the reference, we use a bimodal bridge dataset (*X*_bridge_*, Z*_bridge_) similar to the approach of Hao et al. [16]. The bimodal bridge shares omics features with both the reference and the query as an intermediate. To achieve this, we first compute the annotation *Ŷ*_bridge_ based on *X*_bridge_ and the regression coefficient matrix *B* from PLS(*X*_ref_ *, Y*_ref_ ) (see point 1 above). We then train a second PLS model PLS(*Z*_bridge_*, Ŷ*_bridge_) to obtain the second regression coefficient matrix *B*_bridge_ of size (*Q × K*). The predicted phenotype of the query *Ŷ*_query_ is then calculated based on *Z*_query_ and *B*_bridge_. We illustrate this analysis in the scATAC-seq case study in Section 3.3.

In all cases above, the main output of Φ-Space is the annotation of the *M* query cells on a continuous scale, denoted *Ŷ*_query_ of size (*M × K*), which is calculated either based on *X*_query_ (within-omics), *Z*_query_ (cross-omics) or both *X*_query_ and *Z*_query_ (multi-omics). We refer to *Ŷ*_query_ as the *phenotype space embedding* of the query data.

### 2.2 Downstream analyses

The phenotype space embedding *Ŷ*_query_ contains rich phenotypic information, which we then leverage in the following downstream analyses.

#### Phenotype space reference mapping (Fig. 1E1)

Mapping the query data to a reduced-dimensional space of the reference is a common way for visualising and comparing the reference and query samples in the same low-dimensional space. However, due to the platform effects between reference and query, reference mapping based on omics features is not always straightforward [14, 15, 21]. This problem can be solved by doing reference mapping based on the phenotype space embeddings of reference and query samples. To achieve this, once we have computed *Ŷ*_ref_ from the reference (Fig. 1B2), we obtain the principal components (PCs) of *Ŷ*_ref_ and then map *Ŷ*_query_ to these PCs (Fig. 1E1). In the DC case study (Section 3.1), we will demonstrate that our phenotype space reference mapping effectively overcomes the discrepancy between the reference and the query datasets. This discrepancy can be caused by different sequencing platforms (e.g. bulk vs single-cell) or even different omics types (e.g. scRNA-seq vs scATAC-seq).

#### Phenotype space marker selection (Fig. 1E2)

Conventional marker selection identifies omics features, such as genes, that distinguish particular groups of samples. Analogously, given the phenotype space embedding *Ŷ*_query_ and some known grouping of the query cells (e.g. donor’s disease condition), we can identify phenotypic markers of different groups of query cells. In the CITE-seq case study (Section 3.2), we show how we can identify cell type and disease severity signatures of COVID-19 patients with different pre-existing autoimmune conditions.

#### Phenotype space clustering (Fig. 1E3)

The phenotype space embedding *Ŷ*_query_ of the query data *X*_query_ (resp. *Z*_query_ for cross-omics annotation) is robust against possible batch effects in *X*_query_ (resp. *Z*_query_), as we demonstrate in the CITE-seq and scATAC-seq case studies (Sections 3.2 and 3.3). This is because *Ŷ*_query_ is derived from the PLS regression model learned from the reference rather than the query, and hence is agnostic of the batch effects in the query omics space. This means that *Ŷ*_query_ can be interpreted as a dimension reduction technique that preserves biological sources of variation, and is suitable for identifying biologically meaningful clusters of cells. A similar approach has been used by Aran et al. [12] who identified a novel subtype of macrophage in their scRNA-seq data. Our phenotype space clustering is a generalisation of their approach to cluster other omics types. See Section 3.3 for an extension to scATAC-seq data.

#### Hard classification (Fig. 1E4)

Similar to conventional cell type annotation methods such as Seurat V3 [9] and SingleR [12], we can also derive predicted cell type labels for query cells. To achieve this, we assign to each query cell the cell type label with the highest predicted scores in each row of *Ŷ*_query_, akin to hard classification. We demonstrate in our benchmark studies that Φ-Space is able to achieve accurate cell type annotation in the scATAC-seq case study (Section 3.3).

### 2.3 Case studies

We analysed three biological case studies to showcase the versatile usability of our Φ-Space framework. These case studies are summarised in Table 1.

**Table 1.** Summary of case studies. *N* denotes the number of samples (whether bulk or single cell). Phenotypes refer to reference phenotypes that are transferred to query, where *ϕ* denotes the number of phenotype categories. Featured analyses include types of Φ-Space analyses applied to each case study (see Methods Section 2). CITE-seq: cellular indexing of transcriptomes and epitopes by sequencing; scATAC-seq: single-cell assay for transposase-accessible chromatin with; PBMC: peripheral blood mononuclear cell; BMMC: bone marrow mononuclear cell.

| Case study | Reference | Phenotypes | Query | Featured analyses | Challenges |
| --- | --- | --- | --- | --- | --- |
| DC: Developing identity of induced dendritic cells | Stemformatics human DC atlas [23], $N = 341$ bulk RNA-seq samples of DC and monocytes from 14 studies | Cell types ($\phi = 7$) & cell culture methods ($\phi = 4$) | scRNA-seq of $N = 33,525$ *in vitro* induced DCs and some control cell types | 1) Within-omics annotation;<br>2) phenotype space reference mapping;<br>3) Phenotype space marker selection | 1) Strong batch effects in bulk reference;<br>2) Large differences in sequencing depth and sparsity between bulk and single-cell samples;<br>3) Developing cell identities of induced DCs |
| CITE-seq: Comparing immune cell composition and severity of COVID-19 patients with different pre-existing autoimmune conditions | CITE-seq PBMC atlas of COVID-19 patients with different severity [27], $N = 64,733$ cells from 130 donors | Broad ($\phi = 18$) and fine ($\phi = 51$) cell types & COVID-19 severity ($\phi = 9$) | CITE-seq PBMC data from COVID-19 patients with pre-existing autoimmune conditions [25], $N = 97,499$ from 22 patients | 1) Multi-omics annotation;<br>2) phenotype space marker selection | 1) Integration of two modalities (RNA & protein) in both reference and query;<br>2) Quantitative characterisation of COVID-19 severity of autoimmune patients |
| scATAC-seq: Cell type classification within or between omics types | One out of 13 batches of a BMMC scRNA-seq dataset; we use a cross-validation procedure as described in Methods Section 2.3 | Cell types ($\phi = 27$) | BMMC scRNA-seq (within-omics annotation) or BMMC scATAC-seq (cross-omics annotation); as described in Methods Section 2.3 | 1) Cross-omics annotation;<br>2) hard classification;<br>3) Phenotype space clustering | 1) Differentiating cell types in BMMC;<br>2) Batch effects between reference and query;<br>3) Accurate cell type transfer from scRNA-seq to scATAC-seq |

#### Case study 1: Dendritic Cells (DC)

We show Φ-Space’s ability to characterise developing cell identity by projecting scRNA-seq data from Rosa et al. [22] to a bulk RNA-seq reference atlas generated by Elahi et al. [23]. The query scRNA-seq dataset consisted of 33, 525 *in vitro* induced human dendritic cells (DCs) and some *in vivo* DCs served as control cell types. The bulk reference RNA-seq data consisted of 341 bulk RNA-seq samples of different DC and monocyte subtypes from 14 studies and 10 laboratories. We applied Φ-Space to continuously phenotype the query cells using the 7 cell types and 4 cell culture methods defined in the reference.

#### Case study 2: CITE-seq

We illustrate Φ-Space’s ability to integrate different omics modalities in a common phenotype space. We used cellular indexing of transcriptomes and epitopes by sequencing (CITE-seq) data from Stoeckius et al. [24] as the reference and queried CITE-seq data from Barmada et al. [25]. CITE-seq provides simultaneous measurement of transcripts (RNA) and surface proteins (ADT, antibody-derived tag) at the single-cell level [24]. The reference atlas contained 64,733 peripheral blood mononuclear cells (PBMCs) from 130 donors with different levels of COVID-19 severity, whereas the query contained 97,499 PBMCs from 22 COVID-19 patients with different types of pre-existing autoimmune diseases. We applied Φ-Space to phenotype the query cells using the 18 broad cell types, 51 fine cell types and 9 categories of COVID-19 severity.

#### Case study 3: scATAC-seq

We systematically benchmarked Φ-Space’s ability to transfer cell type labels within or across omics types. We used the bone marrow mononuclear cell (BMMC) 10x multiome dataset from Luecken et al. [26]. This bimodal dataset contains matched RNA and assay for transposase-accessible chromatin with sequencing (ATAC) measurements of 69,249 BMMCs from 13 experimental batches, all from healthy donors. The ATAC measurements are available in two resolutions: peaks (computationally inferred open chromatin regions) and gene activity scores (gene level aggregated peaks). Luecken et al. [26] manually annotated the cells, and we use their annotation as ground truth. We designed two cross-validation (CV) procedures:

- For within-omics annotation, we used only the RNA modality of the 10x multiome dataset for CV. In each of the 13 CV iterations, we used 1 of the 13 batches of annotated scRNA-seq data as the reference and the remaining 12 batches as queries (with ground truth labels hidden).
- For cross-omics annotation, we used both the RNA and ATAC modalities. An additional well annotated BMMC scRNA-seq dataset from a single healthy donor [9] was used as the reference. Then, in each of the 13 CV iterations, we used 1 of the 13 batches of bimodal data as the bridge (Fig. 1D). For the remaining 12 batches, we used only their ATAC modality as scATAC-seq queries (with ground truth labels hidden).

Classification errors, compared to ground truth cell type labels in the query, were then calculated. In the cross-omics annotation case above, since the cell type labels in the reference and those in the multiome data were named according to different conventions, we manually regrouped the labels into common broad cell type labels to make the calculation of classification error possible (see Supplementary Tables S1–S2 for details).

## 3 Results

### 3.1 Continuous characterisation of developing cell states

Conventional cell type annotation methods assign a known cell type label in the reference to each query cell. While suitable for identifying well characterised and typical cell types, these methods are under-powered in characterising transitional or atypical cell identities [8, 9]. In addition, these methods assume each sample or cell has a unique cell type label and thus fail in considering multiple layers of phenotypes. Also, in a typical computational workflow [8, 28], cell type annotation is deployed as a separate step, independent from batch effect removal. This may lead to suboptimal annotation results since phenotypic variation is often confounded with experimental batches (e.g. certain cell types or disease conditions only come from certain batches). Thus a complete removal of batch effects may entail partial loss of between-phenotype heterogeneity.

To address these problems, Φ-Space provides a holistic approach where batch effects removal is tailored for cell type annotation. Instead of assigning a unique cell type label to each query cell, Φ-Space enables a joint modelling of multiple layers of phenotypes. In addition, the Φ-Space continuous phenotyping results are effective for characterising transitional cell states. We illustrate these appealing characteristics of Φ-Space using our first case study below. Our analyses described below suggest that Φ-Space can remove strong batch effects in the reference atlas of Elahi et al. [23], while preserving enough biology to characterise transitional cell states of the induced dentritic cells (DCs) of Rosa et al. [22].

#### Φ-Space selects genes to remove unwanted variation in bulk data

Since the DC dataset from Elahi et al. [23] contains a large number of 341 samples generated by multiple cell culture methods, it can serve as a reference for benchmarking new *in vitro* models of DC biology. In particular, it is suitable for testing if the reprogramming method of Rosa et al. [22] has successfully cultured induced DCs. However, in the original bulk reference dataset with 16,562 genes, the batch effects caused by the sequencing platform dominates the variation (Fig. 2A). To remove the strong platform effects, Elahi et al. [23] applied a feature selection approach introduced by Angel et al. [14] to filter out genes whose variations were mainly explained by the platform, resulting in a total of 2,416 genes were kept and the platform effects were much alleviated (Fig. 2B). However, since the platform effects were confounded with the sample source, the differences between, say, *in vitro* and *in vivo* samples were also blurred in the PC space. In contrast, Φ-Space selected a smaller subset of 1,822 features, which retained some platform effects but, as a result, also preserved a better separation of sample sources (Fig. 2C).

**Fig. 2.**
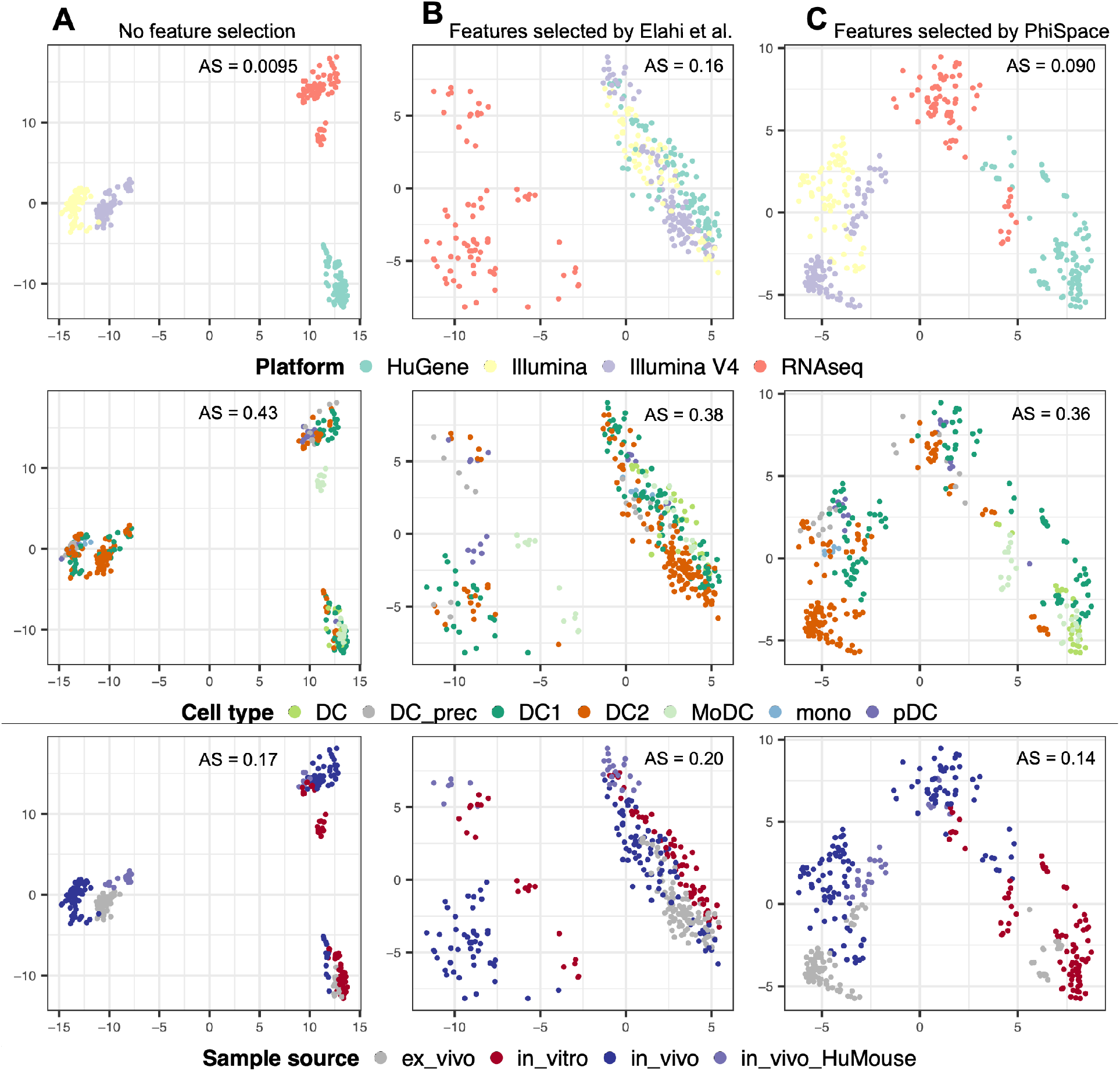
Benchmarking feature selection for building reference atlas. Each column shows all bulk reference samples from Elahi et al. [23] viewed from PC1 and PC2 computed using: **A**) all features, **B**) features selected by Elahi et al. and **C**) features selected by Φ-Space. The samples are coloured by platform, cell type and sample source. AS: alignment score computed using 20 PCs (see Supplementary Methods Section S2.2); larger AS implies better mixing (equivalently poorer separation) of samples from different conditions. Without any gene filtering, the platform effects were the strongest (AS=0.0095) and the separation of cell types was the poorest (AS=0.43); the feature selection of Elahi et al. [23] significantly removed platform effects (AS=0.16, best mixing of batches) and had a better separation of cell types (AS=0.38), but it led to the poorest separation of sample source (AS=0.20); Φ-Space had the best separation of cell types (AS=0.36) and sample source (AS=0.14) by preserving some platform effects that confounds the sample source (AS=0.090).

#### Φ-Space phenotype space reference mapping reveals developing cell identity

Even though Fig. 2C provides a suitable reference atlas for DC biology, mapping scRNA-seq to this atlas remains challenging. This is due to the technical differences between bulk and single-cell RNA-seq data, including the much higher proportion of zero counts in the single-cell query compared to the bulk reference. A naive PCA-based reference mapping (Supplementary Methods Section S2.3) resulted in visualisation that is difficult to interpret (Fig. 3A1). Deng et al. [15] tackled this problem by proposing an imputation method to reduce the sparsity of the scRNA-seq query. However, Sincast imputation did not significantly improve the interpretability of reference mapping results (Fig. 3A2): while the *in vivo* DC subtypes (DC1, DC2 and pDC) showed improved alignment with their bulk counterparts, the induced DCs (Day3, Day6 and Day9) remained misaligned with bulk samples. This result suggested that imputation might not be sufficient for bridging the gaps between bulk and single-cell RNA-seq data in this case.

**Fig. 3.**
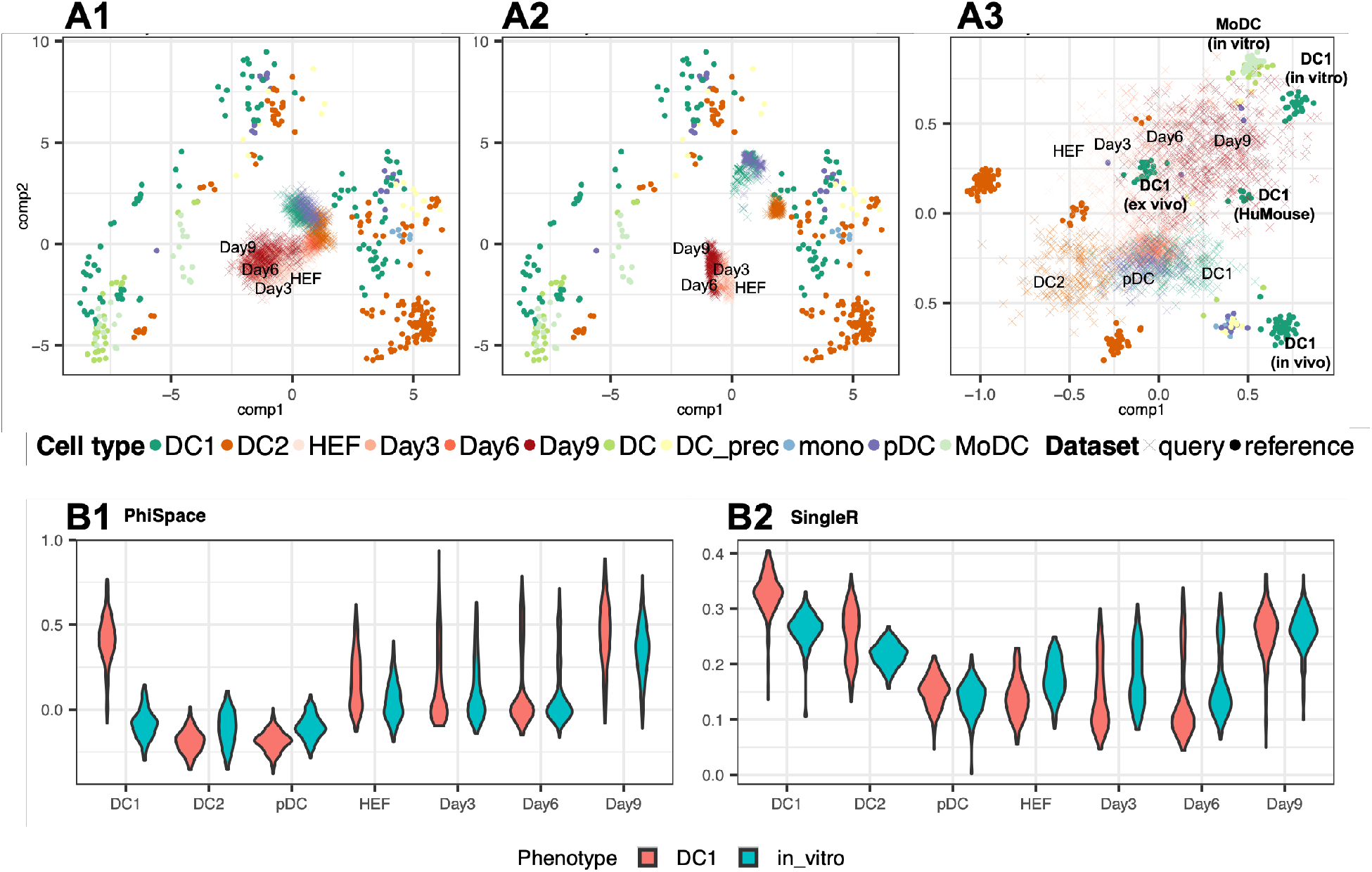
Characterisation of developing dendritic cell (DC) identities. **A**) Benchmarking reference mapping in gene space and in phenotype space, showing bulk reference samples (*•*) from Elahi et al. [23] and query single cells (*×*) from Rosa et al. [22]. HEFs: human esophagus fibroblasts; Day3, Day6 and Day9: induced DCs, i.e. HEFs after 3, 6 and 9 days’ reprogramming towards DCs. **A1**) We first built bulk reference atlas by conducting a PCA of the reference gene expression using genes selected by Φ-Space (depicted in Fig. 2C) and the query gene expression data were mapped to PC1 and PC2 using the bulk reference PCA loadings. The query cells were concentrated around the origin (0, 0) due to the high sparsity of the scRNA-seq gene expression matrix. **A2**) We then mapped scRNA-seq data imputed using Sincast [15] to the same bulk atlas. The induced DCs were still misaligned with the bulk samples. **A3**) Lastly we built bulk reference atlas by conducting a PCA of phenotype space embeddings of the bulk reference and the phenotype space embedding of query cells were then projected onto PC1 and PC2. A convergence of reprogrammed HEFs towards *in vitro* type-1 conventional dendritic cell (DC1) could be observed. **B**) Benchmarking influence of joint modelling on continuous phenotyping: **B1**) Φ-Space annotation, where a gradual increase of the DC1 and *in vitro* identities during 9 days of reprogramming could be observed, and where the *in vivo* non-DC1 control cell types (DC2 and pDC) showed very low DC1 and *in vitro* scores; **B2**) SingleR scores computed using genes selected by Φ-Space, where a comparable transition of the induced DCs was observable, but the DC2 cells were assigned overly high DC1 scores.

Unlike conventional annotation methods, Φ-Space focuses on the phenotype space rather than the gene expression space. This approach enables us to circumvent the harmonisation of bulk reference and single-cell query. Recall from Methods Section 2.2 that we first compute the PCs of *Ŷ*_ref_ , instead of the gene expression matrix *X*_ref_ , and then map *Ŷ*_query_ to these PCs. Fig. 3A3 shows the phenotype space reference mapping results for the DC case. After 3, 6 and 9 days of *in vitro* reprogramming, the induced DCs converged to the bulk *in vitro* type-1 conventional dendritic cell (DC1) samples in the reference. In contrast, the DC1s in the query, used by Rosa et al. [22] as the control, tended to be closer to their bulk counterparts compared to all other cells. Overall we observed that the *in vivo* and induced DCs tended towards two different directions defined by the reference phenotypes. This was confirmed by the heatmap representation of the phenotype space embedding of query cells (Supplementary Fig. S1).

#### Φ-Space can jointly models two layers of phenotypes

In addition to removing unwanted source of variation, Φ-Space’s joint modelling of cell type and sample source yielded more interpretable continuous phenotyping results (Fig. 3B). This can be seen by comparing our results to SinlgeR [12], which also provides continuous phenotyping but can only separately model cell type and sample source. From the Φ-Space results (Fig. 3B1) we observed a gradual increase of both DC1 and *in vitro* scores of induced DCs during the reprogramming, and a clear distinction of the three control cell types DC1, DC2 and pDC (all *in vivo*). From the SingleR results (Fig. 3B2), we observed a comparable transition from HEF to Day 9 induced DCs. However, the SingleR results suggested that the *in vivo* DC1 and DC2 in the query had strong *in vitro* DC1 identity, which was not biologically sensible, suggesting that SingleR was underpowered to jointly model the phenotypic variation pertaining to cell type and sample source.

#### Summary

Through the DC case study, we illustrated how Φ-Space provides a streamlined way for mapping scRNA-seq queries to bulk reference atlases and reveal developing cell identities. We showed that Φ-Space selected biologically meaningful genes to preserve the phenotypic variations in a heterogeneous collection of bulk datasets. To visualise bulk samples and single cells side by side, Φ-Space performs reference mapping using the phenotype space embeddings of reference and query rather than the gene expression. By doing so, our approach does not require explicit harmonisation of these two types of data, such as imputation. In addition, Φ-Space successfully modelled two different layers of cell phenotypes, cell type and sample source, yielding better results compared to modelling them independently. In terms of biological insights, our analyses described above confirmed the claim of Rosa et al. [22] that their experimental method has reprogrammed HFEs to induced DCs with DC1-like transcriptional profile.

### 3.2 Single-cell multi-omics integration in phenotype space

With the maturation of single-cell multi-omic sequencing, single-cell reference atlases consisting of multimodal datasets, e.g. CITE-seq atlases, are becoming more common [24, 27, 29]. Since both the RNA and the ADT modalities of CITE-seq are useful in characterising the cell identities related to immune responses, several methods focusing on integrating these two modalities have been proposed [30–32]. However, a method for jointly querying multimodal data against multimodal references is still lacking. We provide a solution to this problem based on the Φ-Space multi-reference annotation (described in Methods Section 2.2) as follows. First we trained a PLS model using the RNA features for predicting both COVID-19 severity and 51 fine immune cell types. Then we trained another PLS model using the ADT features only for predicting the severity and 18 broad cell types, since the number of ADT features (192) is under-powered for predicting the large number of fine cell types. We then concatenated the predicted phenotype scores for the query cells derived using their RNA and ADT features. Our phenotype space analysis provided some fresh insights to the complex interactions between disease conditions and cell type compositions.

#### Phenotype space embeddings provide biology-preserving integration across batches and modalities

Both the RNA and ADT modalities of the query data showed significant batch effects (Fig. 4A–B). Barmada et al. [25] used totalVI [30], a state-of-the-art integration method for CITE-seq based on variational autoencoders, to integrate the two modalities while removing batch effects. However, this complete removal of batch effects also blurred the boundaries between cell types and rendered cells from different disease conditions no longer distinguishable (Fig. 4C). In contrast, we computed two versions of phenotype space embeddings from either Φ-Space or Seurat V3 (Fig. 4D–E respectively); see Supplementary Methods Section S2.4 for how we applied Seurat V3 from Stuart et al. [9]. Both figures showed well-separated cell types while preserving the differences between disease types. Notably, no batch effects removal step was needed to achieve this separation. This result illustrated the superiority of the phenotype space embedding in preserving complex phenotypic variations and integrating query modalities in the phenotype space.

**Fig. 4.**
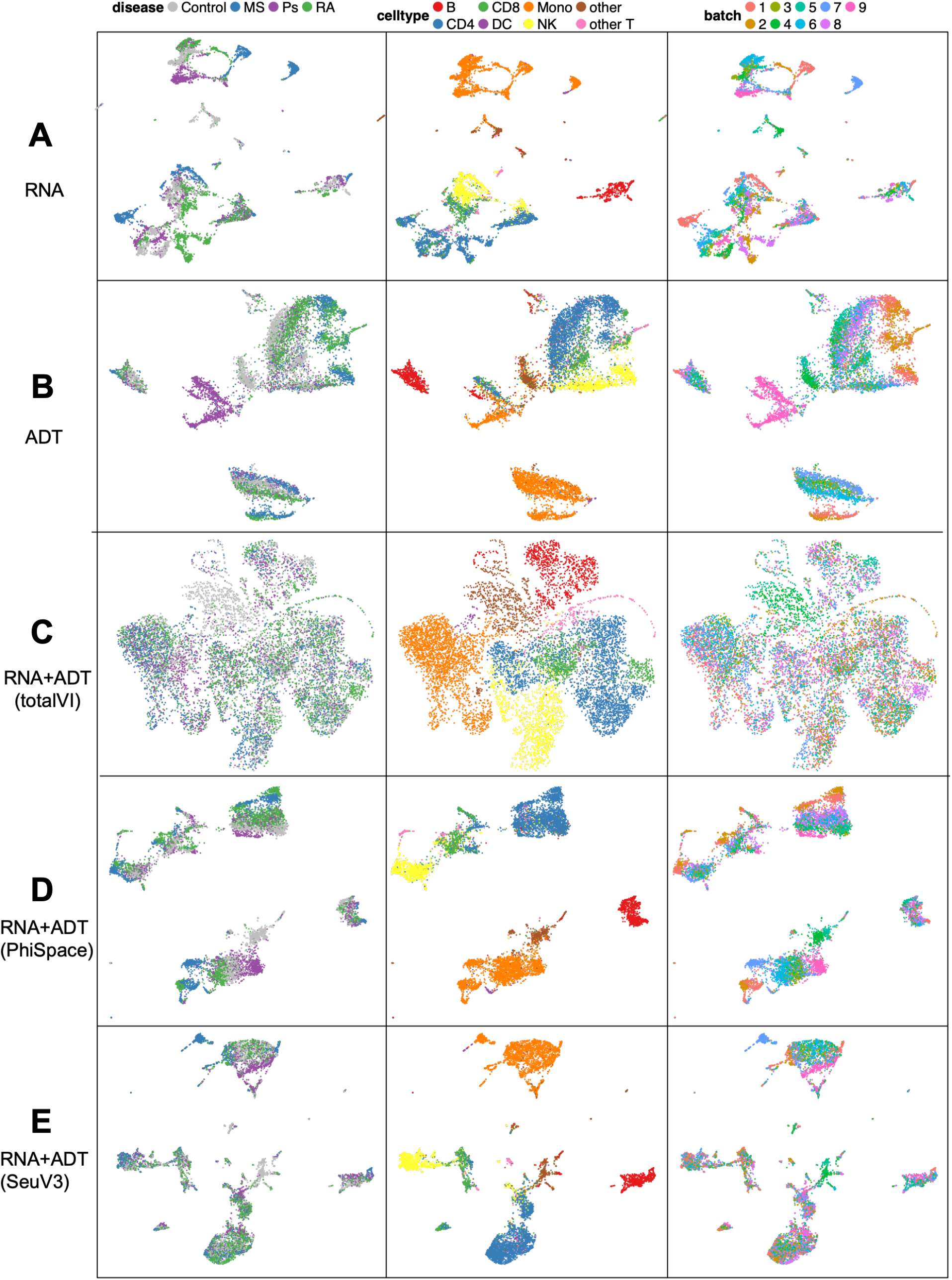
Different representations of query cells in the CITE-seq case study. Uniform manifold approximation and projection plots (UMAPs) based on: **A**) PCs of RNA features, **B**) PCs of ADT features, **C**) 64 latent variables from totalVI [30], computed using both RNA and ADT features, **D**) 64 PCs of the 87-dimensional Φ-Space phenotype space embedding, computed using both RNA and ADT modalities and **E**) 64 PCs of the 87-dimensional Seurat V3 phenotype space embedding, computed using both RNA and ADT modalities. We obtained the totalVI UMAP results from Barmada et al. [25], who set the number of latent variables to be 64. To make our results comparable to the totalVI results, we reduced the dimension of the phenotype space embedding to 64 by PCA. UMAPs of the RNA and ADT modalities showed significant batch effects. TotalVI completely removed the batch effects, but also removed the difference between disease conditions. In contrast, the phenotype space embeddings obtained by Φ-Space and Seurat V3 achieved a better balance between removing batch effects and retaining the difference between disease conditions.

#### Φ-Space characterises complex interactions between cell types and disease conditions

Our phenotype space approach of multi-omics integration also provides a new way of quantifying the interaction between cell types and biological conditions (e.g. disease conditions, age groups). Conventional analyses typically first annotate the single cells, then calculate cell type proportions under each biological condition [27, 33, 34]. However, this approach is very sensitive to the way cell types are defined, which hinders the direct quantitative comparison of cell type compositions derived in different studies. With Φ-Space, we solve this problem by analysing both the reference and query disease conditions in the same phenotype space, so that cell type compositions are directly comparable. The Φ-Space predicted cell type scores differentiated the enrichment or depletion of cell types according to disease conditions (Fig. 5A), in agreement with Stephenson et al. [27] and Barmada et al. [25] (discussed below). Thus we showed that with Φ-Space we can compare cell type compositions across biological conditions and studies. Of note, we could use Seurat V3 scores to plot similar heatmaps (Supplementary Fig. S2). However, the plots were not interpretable due to the overly high proportion of zeros in the scores.

**Fig. 5.**
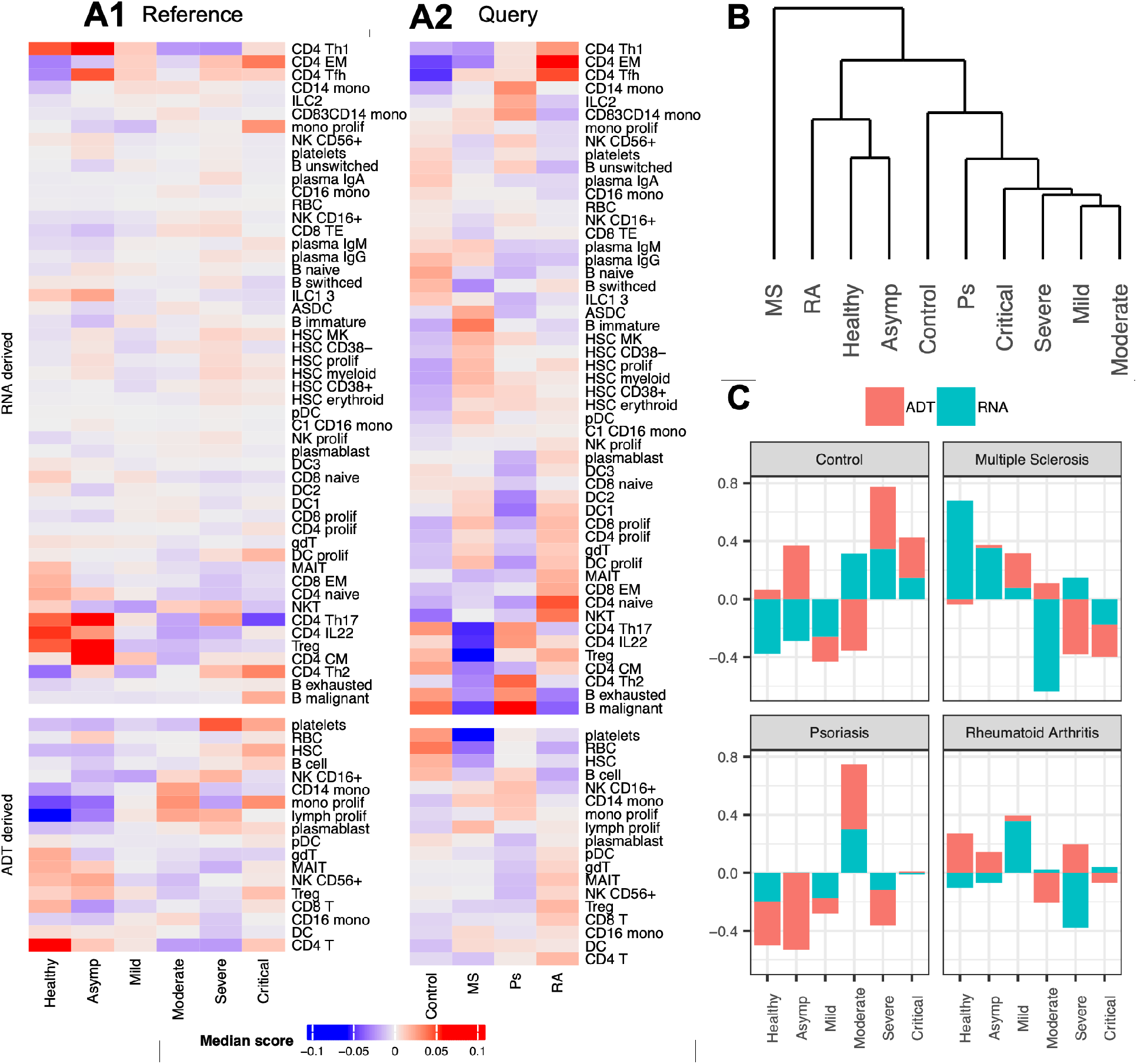
Phenotype space marker selection for CITE-seq data. **A**) Median Φ-Space cell type scores under different disease conditions: **A1**) scores of reference cells, where columns are COVID-19 severity (Asymp: asymptomatic); **A2**) scores of query cells, where columns are types of pre-existing autoimmune diseases (Control: severe COVID-19 patients without pre-existing autoimmune diseases; MS: multiple sclerosis; Ps: psoriasis; RA: rheumatoid arthritis). **B**) Hierarchical clustering of disease conditions in A according to their correlation with cell types. **C**) PLS regression coefficients of COVID-19 severity for predicting autoimmune disease types, where red (or teal) bar corresponds to ADT (or RNA) derived scores. We observed a heterogeneity of COVID-19 patients with different types of pre-existing autoimmune diseases, in terms of cell type composition and disease severity.

Based on the ADT-derived results in severe COVID-19 patients compared to healthy donors, we observed an increased presence of B cells, platelets, plasmablasts, hematopoietic stem cells (HSCs), proliferating monocytes and proliferating lymphocytes (Fig. 5A1). In addition, based on the RNA derived scores, we observed the enrichment of type 1 helper T cells (CD4 Th1), IL22 expressing CD4 T cells (CD4 IL22) and regulatory T cells (Treg) in asymptomatic patients. Both observations are consistent with findings in Stephenson et al. [27]. Moreover, consistent with Barmada et al. [25], we observed in the query data (Fig. 5A2) an overall lack of phenotypic signatures characterising severity, such as HSCs, platelets and plasmablasts, in COVID-19 patients with autoimmune diseases, compared to the severe COVID-19 patients in the control group. Furthermore, Barmada et al. [25] identified an enrichment of type 2 helper T cells (Th2) in psoriasis (Ps) patients and the enrichment of CD4 memory T cells in rheumatoid arthritis (RA) patients, which were confirmed in Fig. 5A2.

In addition to these known findings, our Φ-Space analysis provided a much more nuanced characterisation of the heterogeneity of autoimmune disease types. For example, compared to psoriasis (Ps), both rheumatoid arthritis (RA) and multiple sclerosis (MS) patients tended to display stronger mature DC (DC1, DC2 and DC3) presence, a feature characterising healthy donors (Fig. 5A1; see also Stephenson et al. [27]). In addition, Ps patients were characterised by an enrichment of the exhausted and malignant B cells, which were both indicators for greater COVID-19 severity [27]. These additional findings illustrated the heterogeneity of the immune landscape of COVID-19 patients with different types of autoimmune diseases. This heterogeneity was further illustrated by a hierarchical clustering analysis of cell type composition between disease conditions in Fig. 5B. In light of the relatively small sample size involved in Barmada et al. [25], more samples from COVID-19 patients with autoimmune diseases need to be collected for a thorough investigation of this heterogeneity.

#### Φ-Space highlights disease severity markers of autoimmune conditions

To directly characterise disease severity of COVID-19 patients with autoimmune disease, we conducted a phenotype space marker selection. We used the query cells’ Φ-Space embeddings to predict their autoimmune disease types (Fig. 5C). This allowed us to directly assess how cell type enrichment and COVID-19 severity discriminate autoimmune disease types (only COVID-19 severity was visualised). As expected, the phenotypic variation of the control group consisting of severe COVID-19 patients without any autoimmune disease were best predicted by severe and critical COVID-19 conditions. In contrast, MS and RA were better predicted by healthy, asymptomatic and mild conditions. Finally, Ps was better predicted by the moderate condition. These findings were consistent with our findings shown on Fig. 5A–B. These figures illustrated from different angles the heterogeneity of the immune landscapes of COVID-19 patients with different pre-existing autoimmune diseases.

#### Summary

Through the CITE-seq case study, we demonstrated how a simple concatenation of phenotype space embeddings derived using different omics modalities leads to integrative analyses of complex interactions of cell phenotypes. As a generic modelling strategy, our multi-omics annotation can also be applied using soft classification methods other than PLS, such as Seurat V3. Compared to a direct integration of the omics features, our phenotype space approach achieved a better balance between removing batch effects and preserving difference between disease conditions. This enables us to conduct insightful phenotype space marker analyses, where we could directly compare cell type compositions and disease conditions from different studies. In particular, our approach provides a fully quantitative alternative to the largely qualitative approach used in Barmada et al. [25].

### 3.3 Flexible cell type transfer within and across omics

So far, we have described the ability of Φ-Space to perform soft classification. Here we show that Φ-Space also provides high quality hard classification for both within- and across-omics cell type transfer. After within- or across-omics annotation of the query data, we discretised query cells’ phenotype space embeddings to obtain their predicted cell type labels (see Methods Section 2). We designed two cross-validation (CV) procedures to benchmark the accuracy of Φ-Space against several state-of-art methods in both within- and across-omics label transfer (see summary of case studies in Section 2.3). We considered the complex batch effects in the dataset when designing these CV procedures: they mimic the realistic scenario where the reference and query datasets are generated under different experimental conditions [6, 10].

The balanced classification errors were then calculated based on the ground truth cell type labels (Supplementary Methods Section S2.5). Overall, Φ-Space had very similar classification errors compared to the state-of-the-art Seurat V3 and scANVI, while SingleR led to a much higher error rate (Fig. 6A1).

**Fig. 6.**
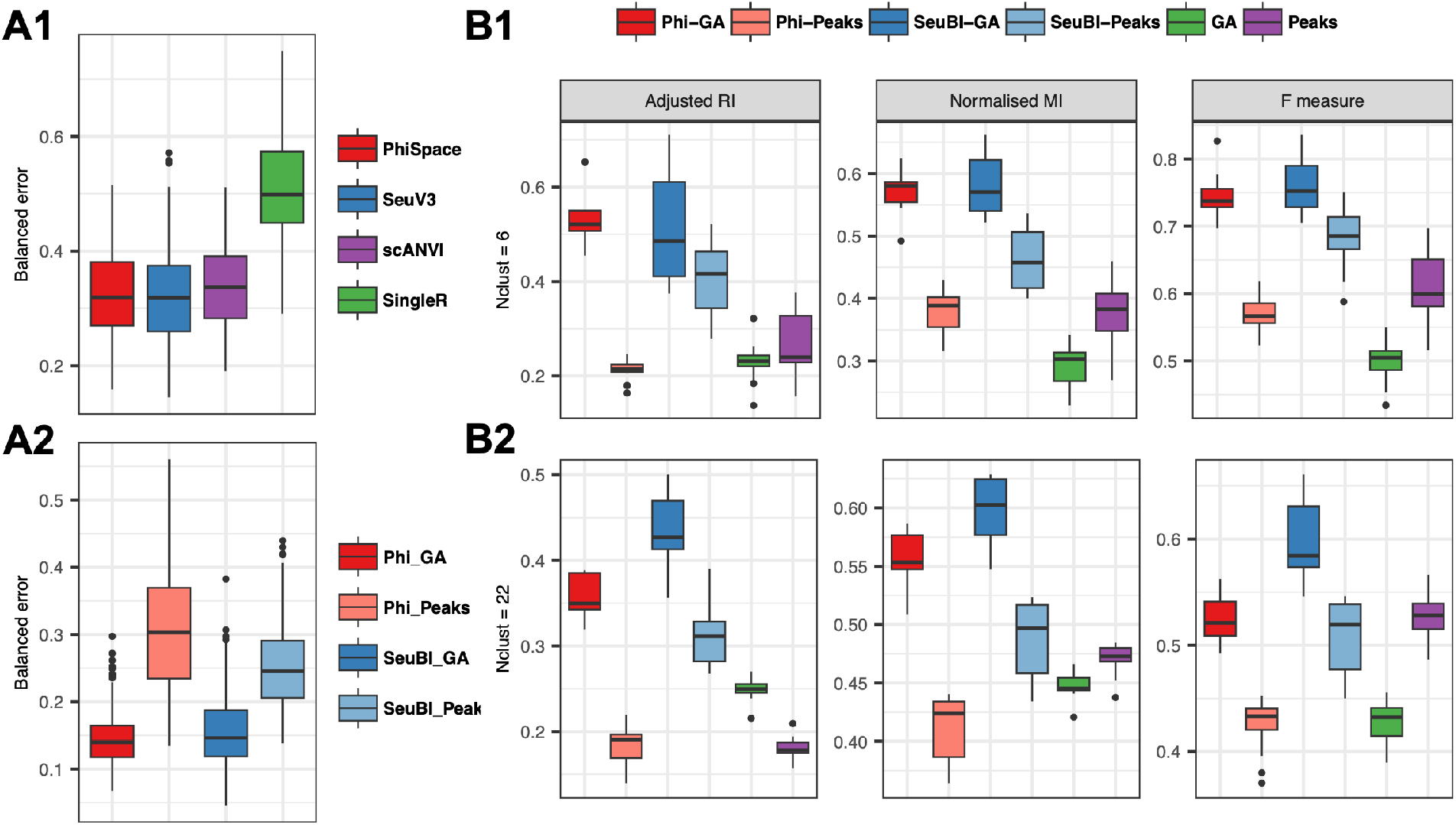
Benchmarking hard classification and phenotype space clustering performances. **A**) Balanced classification errors for: **A1**) within-omics annotation, where we annotated scRNA-seq query using scRNA-seq reference by Φ-Space (Phi), Seurat reference mapping (SeuRefMap), scANVI and SingleR; **A2**) cross-omics annotation, where we annotated scATAC-seq query using scRNA-seq reference and bimodal bridges by Φ-Space and Seurat bridge integration (SeuBI). Since ATAC measurements can be represented as either peaks or gene activity (GA) scores, each of Phi and SeuBI had its ‘Peaks’ and ‘GA’ versions. **B**) Three metrics (adjusted Rand index, normalised mutual information and F measure) for evaluating k-means clustering based on phenotype space embeddings generated by Phi-GA, Phi-Peaks, SeuBI-GA and SeuBI-Peaks, and k-means clustering based on PCs of either GA scores or peaks. The number of k-means clusters were set to be: **B1**) 6, which is the number of ground truth major cell types; or **B2**) 22, which is the number of fine cell types. **C**) UMAPs of **C1**) GA scores and **C2**) Φ-Space embeddings of 8 selected batches of scATAC-seq queries, using the scRNA-seq dataset bmcite [9] as reference and batch s4d8 as bridge. Φ-Space showed performances comparable to state-of-the-art methods in both within- and cross-omics annotation. In terms of clustering, Φ-Space was always among the top two best performing methods.

#### Φ-Space accurately transfers cell type labels from scRNA-seq to scATAC-seq datasets

In our second CV procedure, we used both the RNA and ATAC modalities of the 10x multiome dataset to evaluate cross-omics label transfer performances of Φ-Space and Seurat bridge integration (SeuBI) [16]. ATAC measurements can be represented as either peaks or gene activity (GA) scores. Therefore, we evaluated the classification performance of Φ-Space and SeuBI on each of these representations. On GA scores, both Φ-Space and SeuBI had similar classification error rates, with the Φ-Space errors being slightly less variable (Fig. 6A2); on peaks, the classification error rates were noticeably higher than on the GA scores for both methods, suggesting that GA scores rather than peaks lead to better performance for cell type transfer.

#### Phenotype space clustering leads to accurate identification of out-of-reference cell types

A limitation of fully supervised cell type transfer is that cell types present in the query are not necessarily defined in the reference (e.g. cancer cells that are not present in healthy reference). Our phenotype space clustering strategy (described in Supplementary Section S1) provides a solution to this problem. We adapted the above CV framework for evaluating clustering, where in each CV iteration we simply concatenated the phenotype space embeddings of query cells and applied k-means clustering (Supplementary Section S2.6). Since SeuBI also provided some predicted cell type scores. To illustrate the advantage of phenotype space clustering rather than the conventional ATAC feature space clustering, we also applied k-means to peaks and GA scores directly (Supplementary Methods Section S2.6). We evaluated all methods’ ability to recover either the 6 broad ground truth cell types or the 22 fine ground truth cell types by specifying *k* = 6 or 22 respectively in each k-means analysis, and used three commonly used metrics to evaluate the clustering results (described in Supplementary Section S2.7).

Φ-Space and SeuBI were the best performing methods, according to all three metrics, when using the GA scores to recover broad cell types (Fig. 6B1). However, the performances of SeuBI on either GA or peaks were much more variable compared to Φ-Space on GA scores, indicating that Φ-Space might be more robust against the quality of reference and bridge datasets. SeuBI, followed by Φ-Space on the GA scores, was the best performing method in recovering fine cell types (Fig. 6B2). Overall, Φ-Space and SeuBI always performed better on the GA scores than the peaks, as confirmed in the classification analyses (Fig. 6A2).

#### Summary

Through the scATAC-seq case study we showed that Φ-Space led to good performances in both within- and cross-omics cell type transfer, comparable to several state-of-the-art methods. Of note, our application of SeuBI on GA scores is novel, as SeuBI was originally designed for utilising peaks [16]. Our benchmarking results suggested that using GA scores (aggregated peaks) might be beneficial for cell type transfer tasks, due to the their higher signal-to-noise ratio compared to peaks.

## 4 Discussion

We developed Φ-Space to address numerous challenges faced by state-of-the-art automated annotation methods, including identifying continuous and out-of-reference cell states, dealing with batch effects in reference, utilising bulk references and including multi-omic references and queries. Φ-Space uses soft classification to phenotype cells on a continuum. We then use the continuous annotation, or phenotype space embedding, to reduce the dimensionality of our data for various downstream analyses.

Through the three biological case studies (DC, CITE-seq and scATAC-seq), we have demonstrated the versatile use of Φ-Space in continuous phenotyping. While our case studies featured some alternative methods, none of these methods can be applied to all modelling tasks described above. In contrast, thanks to the flexibility of PLS regression and our phenotype space modelling strategies, Φ-Space can be applied to a much wider range of tasks, as summarised below.

First, Φ-Space can characterise developing and out-of-reference cell states. In the DC case study, Φ-Space characterised developing cell states of the induced dendritic cells, which were not defined in the reference. In the scATAC-seq case study, via the clustering analysis we showed that Φ-Space better preserved cell types defined in query datasets. This greatly extends the utility of automated cell annotation beyond hard cell type classification.

Second, Φ-Space is robust against batch effects in both reference and query datasets, without requiring additional batch effects correction. This was demonstrated in all three case studies. This greatly simplifies the cell annotation workflow, leading to reduced computational cost.

Third, Φ-Space is flexible enough to be extended to annotation tasks involving multiple omics types. In addition to within-omics annotation (DC case), Φ-Space can also accomplish cross-omics annotation (scATAC-seq case) and multi-omics annotation (CITE-seq case). In particular, our cross-omics annotation approach cannot be replicated by alternative methods such as SingleR or Seurat V3 as we use the bridge dataset to annotate the query, then train a PLS regression model to predict a continuous phenotype matrix – the latter cannot be achieved by these alternative soft classification methods. As single-cell multi-omics data are becoming increasingly common [35], we anticipate that cross-omics and multi-omics annotation will be increasingly useful in biological research.

Fourth, Φ-Space overcomes strong technical differences between reference and query sequencing platforms. In the DC case study, we used bulk reference to annotate scRNA-seq query, utilising a large number of bulk studies. In the scATAC-seq case study, we used scRNA-seq reference to annotate scATAC-seq query. On the one hand, this approach recycles high-quality data from conventional sequencing technologies. On the other hand, emerging sequencing platforms (e.g. the fast evolving spatial transcriptomics [36]) have very different characteristics compared to well established platforms, and we anticipate that Φ-Space will help transfer biological information to these newly generated datasets.

Methodologically speaking, the essence of Φ-Space, namely the phenotype space analysis, is the use of soft classification as dimension reduction. This modelling strategy is greatly under-utilised in computational biology. Our work proved that this modelling strategy leads to very flexible tools for modelling complex phenotypic variations in multi-omics data. In addition, Φ-Space used a particular soft classification method, PLS, a workhorse method deployed in routine analyses of many scientific disciplines [20, 37–39]. However, any soft classification method can be used for Φ-Space type of analyses. This further proves the value of Φ-Space, not just as a versatile computational toolkit for continuously modelling cell phenotypes, but also as a generic modelling strategy.

Φ-Space also has great potential in uncovering complex interactions of different cell phenotypes. This was demonstrated in the CITE-seq case study, where we profiled cellular compositions of COVID-19 patients. Despite a considerable amount of existing work, it remains an open problem how pre-existing autoimmune diseases contribute to COVID-19 severity [40–43]. This is partly due to the complex interaction between the two layers of phenotypic variations, cell types and the donors’ disease conditions. In addition, the query data in the CITE-seq case study from Barmada et al. [25] were particularly challenging to analyse since the disease condition is seriously confounded with experimental batches. Both challenges were effectively dealt with by Φ-Space.

Φ-Space’s flexibility opens many possible future research directions. One limitation of the current work is that we did not make use of bulk multi-omic reference atlases. A potential extension of Φ-Space is the integration of bulk and single-cell multi-omics data. The motivation is that most existing single-cell multi-omics technologies only generate data with two omics types [35]. Hence the abundance of bulk datasets with 3 or more omics types (e.g. TCGA) may mitigate the lack of such single-cell datasets. Another possible extension is to develop Φ-Space into a unified approach for annotating spatial transcriptomics data generated by radically different platforms. We anticipate that the ability of Φ-Space to uncover continuous phenotypic information will facilitate the discovery of spatial phenotypic patterns. We believe this toolkit will empower biological research such as developmental and cancer biology, which feature complex and atypical cell identities that are hard to identify using conventional methods.

## Declarations

### Code availability

The Φ-Space R package and R code for reproducing results in this paper are available at https://github.com/jiadongm/PhiSpace.

### Competing interests

The authors declare they have no competing interests.

## Acknowledgements

We would like to thank Dr Jarny Choi for helpful discussions.

## Funding

JM was supported in part by the Australian Research Council (ARC) Discovery Project DP200102903. KALC was supported in part by the National Health and Medical Research Council (NHMRC) Career Development fellowship (GNT1159458). YD was supported by the Melbourne Research Scholarship.

## Notes

https://github.com/jiadongm/PhiSpace

### Competing Interest Statement

The authors have declared no competing interest.

