## Supplemental material for "Φ-Space: Continuous phenotyping of single-cell multi-omics data"

### S1 Supplementary methods

#### S1 Details of $\Phi$-Space

##### S1.1 Loss function for parameter tuning

In the $\Phi$-Space framework, each experimental unit (bulk sample or single cell) in the reference is allowed to have multiple class labels (e.g. cell type and sample source). To achieve robust and interpretable continuous phenotyping based on soft classification, we use the residual sum of squares (RSS) to tune the parameters in our PLS model. Given the true $\{-1, 1\}$-valued response matrix $Y$ and a continuous predicted version $\hat{Y}$, the RSS criterion is defined as RSS $= \|Y - \hat{Y}\|_F$, where $\|\cdot\|_F$ denotes the Frobenius norm of matrices (sum of squares of all matrix entries). An increasing number of recent theoretical and empirical machine learning works have suggested that training classification models using RSS tend to yield robust and generalisable results for complex data [1–3].

##### S1.2 PLS component and feature selection

For selecting the number of PLS components `ncomp`, we compute a 5-fold cross-validation (CV) version of the RSS criterion, where we split the reference data into training and validation folds. For atlas-scale single-cell references, computing CV is time consuming. Thus, we first use the rule-of-thumb `ncomp` $= K$ for choosing the number of PLS components without any feature selection, as described in Section 2.1. For smaller-scale references such as bulk references, we plot the CV RSS values computed for an equispaced grid of candidate `ncomp` values on the logarithmic scale. Fig S3A shows such a plot for the bulk reference data [4] used in the DC case study in Section 3.1. We chose `ncomp` $= 30$ since further increase of `ncomp` does not significantly decrease the RSS.

After selecting `ncomp`, and if feature selection is desired (e.g. for removing platform effects confounding biology as in our DC case study), we perform variable selection to characterise each of the $K$ phenotypes in the reference as follows. We select the top `nfeat` features that are most useful for predicting that phenotype and then choose the union of these $K$ sets of features as the selected features. We then use only the selected features to compute a PLS model. Here the goal is to improve computational efficiency and model interpretability rather than prediction accuracy. To evaluate the contribution of each feature towards predicting a given phenotype, we extract the regression coefficient matrix $B$ of size $(P \times K)$ from the PLS model computed using all $P$ features and a selected `ncomp` value. Let $\tilde{B}$ denote the version of $B$ where each entry is replaced by its absolute value. Now the $k$-th column of $\tilde{B}$ can be viewed as importance scores for all $P$ features in predicting the $k$-th phenotype. We then choose the top `nfeat` features with the highest importance scores. Hence we have reduced the problem of feature selection to the selection of `nfeat`, which is again done via examining the RSS plot. Fig S3B shows such a plot for the bulk reference in the DC case study. We chose `nfeat` $= 265$ which minimises the RSS value. This gives us a total of 1822 features; see Section 3.1 for more details on an evaluation of these features.

##### S1.3 Normalisation of phenotype space embeddings

Recall that the $\Phi$-Space scores $\hat{Y}_{\text{query}}$ is defined as a normalised version of the raw PLS scores $X_{\text{query}}B$ of size $(M \times K)$ (or $Z_{\text{query}}B_{\text{bridge}}$ for cross-reference transfer). Given a generic matrix $A$ of size $(M \times K)$, we obtain its normalised version $B = \sigma(A)$ by centering each column of $A$ by its median and scaling it by its largest absolute value. This normalisation step effectively improves the prediction for rarer cell types, thus alleviating the so-called 'masking' problem of linear classifiers for data with imbalanced class proportions [5].

#### S2 Details of case studies

##### S2.1 Preprocessing of data

Here we describe how data were processed before applying Φ-Space. See Section 2.3 for a description of data.

**Case study 1: DC.** Since the bulk reference data [4] have been rank normalised, we also applied rank normalisation to the query scRNA-seq data. Rank normalisation was defined in Angel et al. [6] and we implemented it using the `RankTransf` function in our PhiSpace R package.

**Case study 2: CITE-seq.** The RNA modality in both reference and query were normalised using the R package scran [7]. The ADT modality were normalised using central log ratio (CLR) transform, implemented by the `CLRnorm` function in our PhiSpace R package.

**Case study 3: scATAC-seq.** The RNA modality of all 13 batches were normalised, batch by batch, using scran [7]. For the two representation of ATAC modality: peaks were normalised, batch by bach, using term-frequency inverse-document-frequency as implemented by the `RunTFIDF` function in the R package Signac; gene activity scores were normalised, batch by batch, using scran.

##### S2.2 Alignment score

We computed the alignment score defined in Butler et al. [8]. Our R implementation of alignment score was based on the `alignment_score` function in the `PLSDAbatch` package [9].

##### S2.3 PCA-based reference mapping

To obtain Supplementary Fig S1C, we first computed the leading PCs of the bulk reference gene expression matrix, using only the genes selected by Φ-Space (Supplementary Fig S1A3). We then computed the embedding of the query single cells by projecting the scRNA-seq data to the reference PC1 and PC2, using the loadings computed using the bulk reference.

##### S2.4 Implementation of alternative classification methods

In both the CITE-seq and the scATAC-seq case studies (Sections 3.2 and 3.3), we used the R package Seurat V3 [10]. In addition, in the scATAC-seq case study we also implemented SingleR [11] and scANVI [12, 13] using the default tuning parameters used in their vignettes. To make the results more comparable, we used scran [7] as the normalisation method for all three methods.

- Seurat V3. We followed the Seurat vignette `https://satijalab.org/seurat/articles/integration_mapping`. We skipped the normalisation step using `NormalizeData` and instead included scran normalised data as the `data` slot in the Seurat object. We used the predicted cell type scores as the Seurat V3 phenotype space embedding in the CITE-seq case study, and the predicted cell types in the scATAC-seq case study.

- SingleR. We used `trainSingleR` and `classifySingleR` in the R package SingleR with default parameters. Since SingleR does feature selection implicitly, we did not select highly variable genes (HVGs) before training the model.

- scANVI. We used scANVI as implemented in python library scArches [13]. R package reticulate was used to call python functions from R. We followed the scANVI vignette `https://docs.scarches.org/en/latest/scanvi_surgery_pipeline.html`. We used Seurat to select HVGs before training scANVI, so that scANVI used the same HVGs as our implementation of Seurat cell typing above.

In both the scATAC-seq case studies (Section 3.3), for transferring cell types from scRNA-seq reference to scATAC-seq query, we implemented Seurat bridge integration (SeuBI; [14]) by following the SeuBI vignette `https://satijalab.org/seurat/articles/seurat5_integration_bridge`. We used SCTransform as suggested by that vignette.

##### S2.5 Classification errors

We calculated both overall and balanced classification errors to evaluate the label transfer results. Only balanced error were shown in the main manuscript since the overall error led to the same conclusion as in the scATAC-seq case study (Section 3.3). In particular, given predicted cell type labels $\{\hat{l}_1, \ldots, \hat{l}_n\}$ and ground truth cell type labels $\{l_1, \ldots, l_n\}$, where each $l_i \in \{1, \ldots, K\}$ for $i = 1, \ldots, n$. Then the overall classification error is defined by $(1/n) \sum_{i=1}^{n} \mathbf{1}(\hat{l}_i \neq l_i)$, where $\mathbf{1}(\cdot)$ denote the indicator function. The balanced classification error is defined by $(1/K) \sum_{k=1}^{K} \big\{(1/n_k) \sum_{i=1}^{n} \mathbf{1}(l_i = k) \mathbf{1}(\hat{l}_i \neq l_i)\big\}$, where $n_k = \sum_{i=1}^{n} \mathbf{1}(l_1 = k)$ denotes the number of observations from the $k$-th cell type.

##### S2.6 Details of phenotype and omics space clustering

For phenotype space clustering, we concatenated the 22-dimensional phenotype space embeddings of cells from different query batches and applied k-means clustering by using the R function `kmeans` with the `algorithm="Lloyd"`, `iter.max=500` and `nstart=50`. The same k-means implementation was used for omics space clustering below.

For omics space clustering, we concatenate the normalised peaks or gene expressions from query batches and then computed dimension reduction. For peaks, we computed singular value decomposition (SVD) and used the 2nd to the 23rd SVD components as the input to k-means, to match the dimensionality of phenotype space embeddings. Discarding the first SVD component is recommended by the Seurat workflow since this component is usually heavily contaminated by technical noise (`https://satijalab.org/seurat/articles/seurat5_integration_bridge`). For concatenated GA scores, we computed the top 22 PCs, which were then used as input to k-means.

##### S2.7 Metrics for clustering

We computed adjusted Rand index, normalised mutual information using the `clustComp` function in R package aricode [15]. We used our own R implementation of Van Rijsbergen's F measure [16], which is robust version of the commonly used purity metric; see Amigó et al. [17] for the definition of the F measure and its superiority compared to purity.

### S2 Supplementary figures

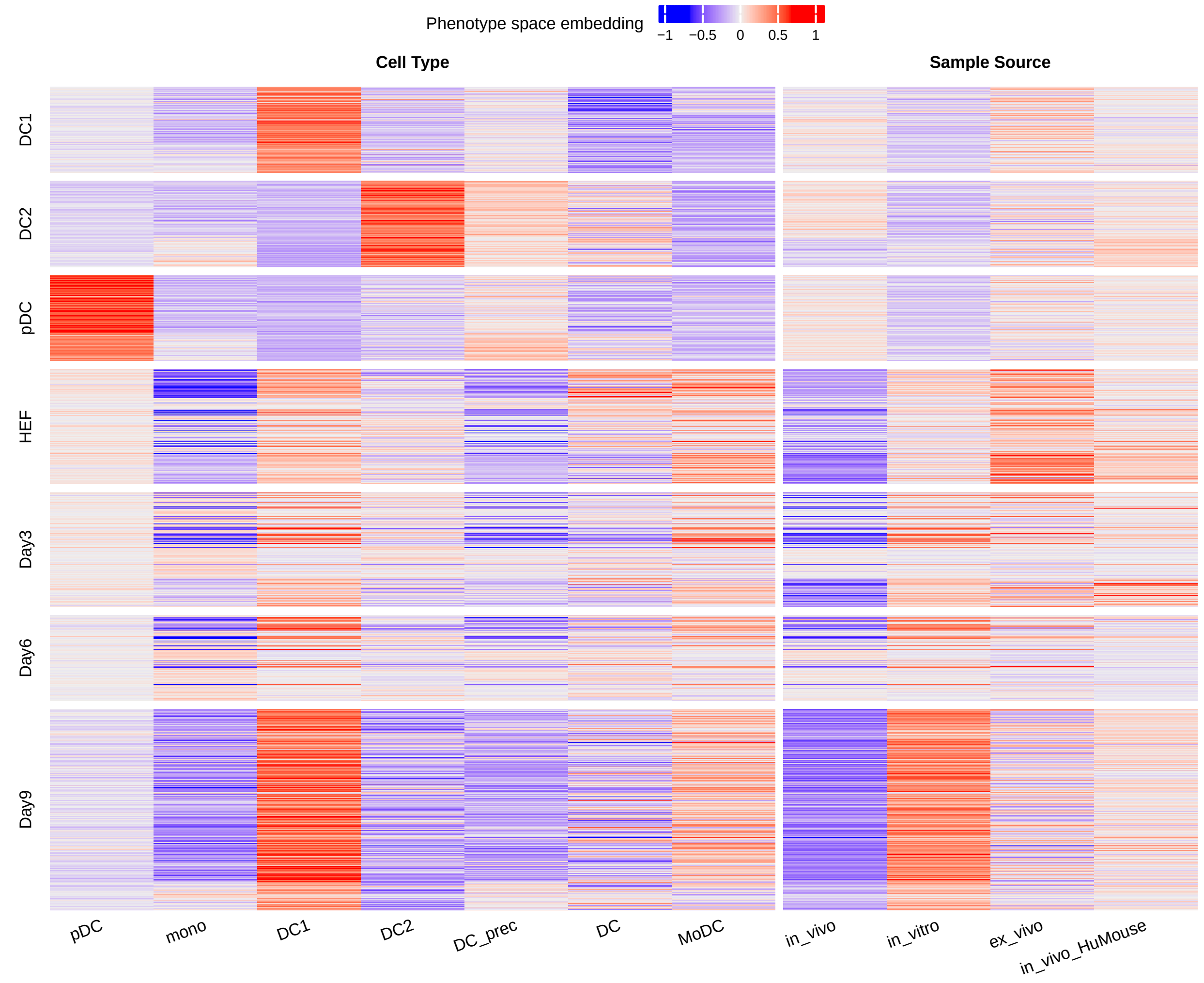


**Fig. S1 DC case study**. Heatmap showing phenotype space embeddings of query cells, where each row is a query cell and each column is a phenotype in the bulk reference. Query cell types include DC1: type 1 conventional dendritic cell (DC); DC2: type 2 conventional DC; pDC: plasmacytoid DC; HEF: human esophagus fibroblast; Day3, Day4, Day9: HEFs after 3, 6 and 9 days of reprogramming towards DC. Reference cell types include, in addition to DC1, DC2 and pDC, mono: monocytes; DC_prec: DC precursor; DC: DC with unknown subtypes; MoDC: monocyte derived DC. The reference sample source in_vivo_HuMouse refers to *in vivo* cells from humanised mice.

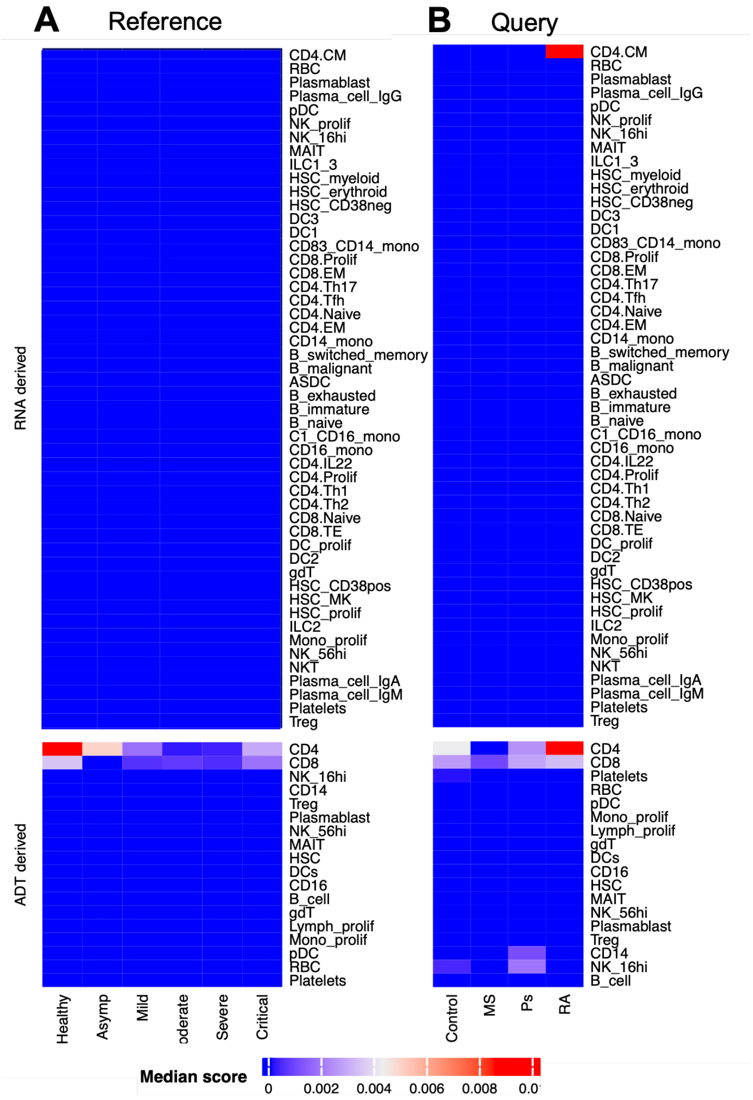


**Fig. S2 CITE-seq case study**. **A–B** are the same as Fig 5A1–A2 except that Seurat V3 predicted cell type scores are used. Most median scores are zero due to the high proportions of zeros in the score matrix.

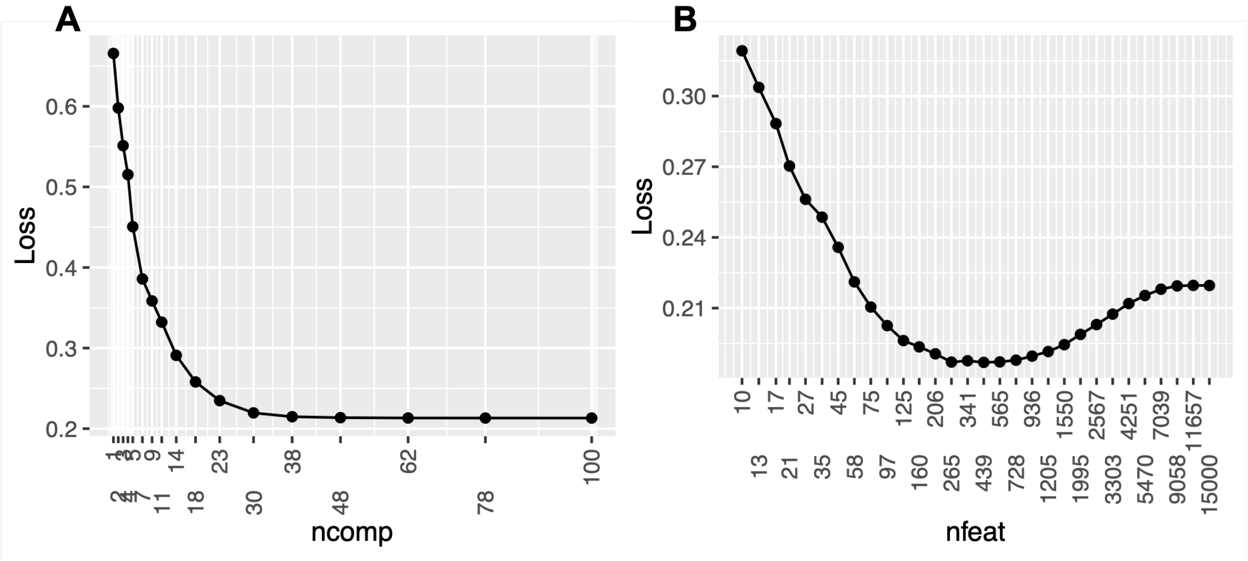


**Fig. S3 Φ-Space parameter tuning for PLS components and feature selection**. Cross-validated residual sum of squares (RSS) defined in Section S1 ($y$-axis) for selecting **A**) `ncomp` and **B**) `nfeat`. Here we can select `ncomp` = 30 and `nfeat` = 265 since they result in minimal RSS values.

### S3 Supplementary tables

**Table S1 Merging fine cell types in the bmcite dataset to broad ones.** We manually merged the fine cell types defined in the bmcite dataset [10] to broad cell types, for benchmark studies on cross-omics annotation (see Section 2.3).

| Fine cell types | Broad cell types |
|---|---|
| CD14 Mono | Mono/DC |
| CD16 Mono | Mono/DC |
| CD4 Memory | T cell |
| CD4 Naive | T cell |
| CD56 bright NK | NK |
| CD8 Effector_1 | T cell |
| CD8 Effector_2 | T cell |
| CD8 Memory_1 | T cell |
| CD8 Memory_2 | T cell |
| CD8 Naive | T cell |
| cDC2 | Mono/DC |
| gdT | T cell |
| GMP | Progenitor cells |
| HSC | Progenitor cells |
| LMPP | Progenitor cells |
| MAIT | T cell |
| Memory B | B cell |
| Naive B | B cell |
| NK | NK |
| pDC | Mono/DC |
| Plasmablast | B cell |
| Prog_B 1 | Progenitor cells |
| Prog_B 2 | Progenitor cells |
| Prog_DC | Progenitor cells |
| Prog_Mk | Progenitor cells |
| Prog_RBC | Progenitor cells |
| Treg | T cell |

**Table S2 Merging fine cell types in the 10x Multiome dataset to broad ones.** We manually merged the fine cell types defined in the 10x Multiome dataset [18] to broad cell types, for benchmark studies on cross-omics annotation (see Section 2.3).

| Fine cell types | Broad cell types |
|---|---|
| B1 B | B cell |
| CD14+ Mono | Mono/DC |
| CD16+ Mono | Mono/DC |
| CD4+ T activated | T cell |
| CD4+ T naive | T cell |
| CD8+ T | T cell |
| CD8+ T naive | T cell |
| cDC2 | Mono/DC |
| Erythroblast | Progenitor cells |
| G/M prog | Progenitor cells |
| HSC | Progenitor cells |
| ID2-hi myeloid prog | Progenitor cells |
| ILC | ILC |
| Lymph prog | Progenitor cells |
| MK/E prog | Progenitor cells |
| Naive CD20+ B | B cell |
| NK | NK |
| Normoblast | Progenitor cells |
| pDC | Mono/DC |
| Plasma cell | B cell |
| Proerythroblast | Progenitor cells |
| Transitional B | B cell |

[2] Like Hui and Mikhail Belkin. Evaluation of neural architectures trained with square loss vs Cross-Entropy in classification tasks. 2020.

[3] Tianyang Hu, Jun Wang, Wenjia Wang, and Zhenguo Li. Understanding square loss in training overparametrized neural network classifiers. 2021.

[4] Zahra Elahi, Paul W Angel, Suzanne K Butcher, Nadia Rajab, Jarny Choi, Yidi Deng, Justine D Mintern, Kristen Radford, and Christine A Wells. The human dendritic cell atlas: An integrated transcriptional tool to study human dendritic cell biology. *J. Immunol.*, 209(12): 2352–2361, 2022.

[5] Trevor Hastie, Robert Tibshirani, and Jerome Friedman. *The Elements of Statistical Learning*. Springer New York, 2009.

[6] Paul W Angel, Nadia Rajab, Yidi Deng, Chris M Pacheco, Tyrone Chen, Kim-Anh Lê Cao, Jarny Choi, and Christine A Wells. A simple, scalable approach to building a cross-platform transcriptome atlas. *PLoS Comput. Biol.*, 16(9), 2020.

[7] Aaron T L Lun, Karsten Bach, and John C Marioni. Pooling across cells to normalize single-cell RNA sequencing data with many zero counts. *Genome Biol.*, 17, 2016.

[18] Malte Luecken, Daniel Burkhardt, Robrecht Cannoodt, Christopher Lance, Aditi Agrawal, Hananeh Aliee, Ann Chen, Louise Deconinck, Angela Detweiler, Alejandro Granados, Shelly Huynh, Laura Isacco, Yang Kim, Dominik Klein, BONY DE KUMAR, Sunil Kuppasani, Heiko Lickert, Aaron McGeever, Joaquin Melgarejo, Honey Mekonen, Maurizio Morri, Michaela Müller, Norma Neff, Sheryl Paul, Bastian Rieck, Kaylie Schneider, Scott Steelman, Michael Sterr, Daniel Treacy, Alexander Tong, Alexandra-Chloe Villani, Guilin Wang, Jia Yan, Ce Zhang, Angela Pisco, Smita Krishnaswamy, Fabian Theis, and Jonathan M Bloom. A sandbox for prediction and integration of dna, rna, and proteins in single cells. In J. Vanschoren and S. Yeung, editors, *Proceedings of the Neural Information Processing Systems Track on Datasets and Benchmarks*, volume 1, 2021.
